# Trackable and scalable LC-MS metabolomics data processing using asari

**DOI:** 10.1101/2022.06.10.495665

**Authors:** Shuzhao Li, Amnah Siddiqa, Maheshwor Thapa, Shujian Zheng

**Affiliations:** Jackson Laboratory for Genomic Medicine, Farmington, CT 06032, USA

## Abstract

Significant challenges still exist in the computational processing of data from LC-MS metabolomic experiments into metabolite features. In this study, we examine the issues of provenance and reproducibility in the current software tools. The inconsistency among these tools is attributed to the deficiencies of mass alignment and controls of feature quality. To address these issues, we have developed a new open-source software tool, asari, for LC-MS metabolomics data processing. Asari is designed with a set of new algorithmic framework and data structures, and all steps are explicitly trackable. Asari compares favorably to other tools in feature detection and quantification. It offers substantial improvement of computational performance over current tools, and is highly scalable.

---

Metabolomics holds the promise to measure and quantify small molecules comprehensively in biological systems. Since the chemistry of these small molecules underlies most aspects of life science, metabolomics is recognized as critical to support missions in biomedical research, including precision medicine and environmental health (Wishart 2016, Barnes 2020, Vermeulen et al, 2020). Since mid-2000s, the experimental platforms of metabolomics have improved significantly, and LC-MS (liquid chromatography coupled mass spectrometry) has become its leading technology.

In LC-MS metabolomics, a sample is scanned by mass spectrometer consecutively during the chromatography, generating a time series of spectra, each containing a list of ions with mass to charge ratio (m/z) and intensity values. The goal of data processing is to report a quantitative value per metabolite feature per sample, which is a proxy of biological concentration. Multiple software tools have been developed for LC-MS metabolomics data processing over the years (Smith et al, 2006, Katajamaa et al, 2006, Pluskal et al, 2010, Du et al, 2020, Yu et al, 2013, Melamud et al, 2010, Rurik et al, 2020, Tsugawa et al, 2020), and the most widely used are XCMS (Smith et al, 2006) and MZmine (Pluskal et al, 2010). They have contributed significantly to the growth of the field, but major design issues have also become apparent over the past decade. As a key –omics technology, the users of metabolomics cannot be limited to chemists, and the data processing has to be trusted by general bioinformatics practitioners. This requires a set of transparent quality metrics and accountability of each major step. In regarding to software design, explicit linking between steps is necessary for automated testing, debugging and continued improvement. The software issues are manifested in the current roadblock of reproducibility.

In global profiling by a typical high-resolution mass spectrometer, studies report several thousand to tens of thousands features. Myers et al (2017) compared XCMS and MZmine 2, and found that half or more of the features were not shared between the tools. Delabriere et al (2021) reported similar level of disagreement between XCMS and OpenMS. It’s common knowledge among users that the results also vary wildly based on parameter settings. Significant community efforts were spent on parameter optimization of XCMS (Uppal et al, 2013, Libiseller et al, 2015, Manier et al, 2018, McLean et al, 2020, Pang et al, 2020, Delabriere et al, 2021). However, these are ad hoc patches, not addressing the fundamentals.

In this study, we present asari, a new open-source software tool for LC-MS metabolomics data processing. Asari is designed with a set of new algorithmic framework and data structures, and all steps are explicitly trackable. The use of mass tracks and composite mass tracks is introduced. They facilitate, along with a set of quality metrics, the better understanding of reproducibility in detecting mass peaks and elution peaks, and correspondence of LC-MS features. Asari offers substantial improvement of computational performance over the previous tools, and is highly scalable.

## Results

### Provenance issues in feature correspondence during LC-MS data preprocessing

Metabolomics today usually employs high-resolution mass spectrometers that are often capable of mass resolution at 5 ppm (part per million) or better. That means, measurement error for a single charged molecule of 150 Dalton is no greater than 0.00075 in m/z values (150 * 5 * 1E-6), and one of 800 Dalton 0.0040 in m/z values (800 * 5 * 1E-6). Mass spectrometry software used to collapse data into m/z bins of nominal or 0.1 amu (atomic mass unit). With the mass resolution in today’s data, binning is no longer a valid approach, and the mass should be reported as precisely as possible to support compound identification. This step is the detection of mass peaks: a group of ions measured on the same molecular species have small random variations, and a consensus m/z value is decided on the group as a mass peak. The detection of mass peaks is now usually taken care by the “centroiding” process; centroided data are then used as input to metabolomics data processing. Centroiding is supported by all major instrument manufacturers. Tools like ThermoRawFileParser (Hulstaert et al, 2020) and msConvert (Adusumilli and Mallick, 2017) are commonly used both for data format conversion and for centroiding.

A feature in LC-MS metabolomics is defined by a unique pair of m/z value and retention time in chromatography. The m/z value of a peak is first determined per sample. When peaks are later aligned cross samples (i.e. correspondence), the m/z values vary slightly in each sample and the algorithm has to ensure grouping the correct peaks and report a consensus m/z value. There are more steps in data processing, while we first focus on m/z alignment.

We generated a LC-MS metabolomic dataset (HZV029) of 184 repeated analyses of a pooled human plasma sample. The same data were processed by XCMS and MZmine, using similar parameters. The two tools produced very different numbers of features, and about 60% of XCMS features are unambiguously matched to MZmine features (**Figure 1A**). Similar levels of disagreement are seen in MZmine using a different algorithm and MS-DIAL (**Tables S1**), and in a different dataset (**Tables S2**).

**Figure 1.**
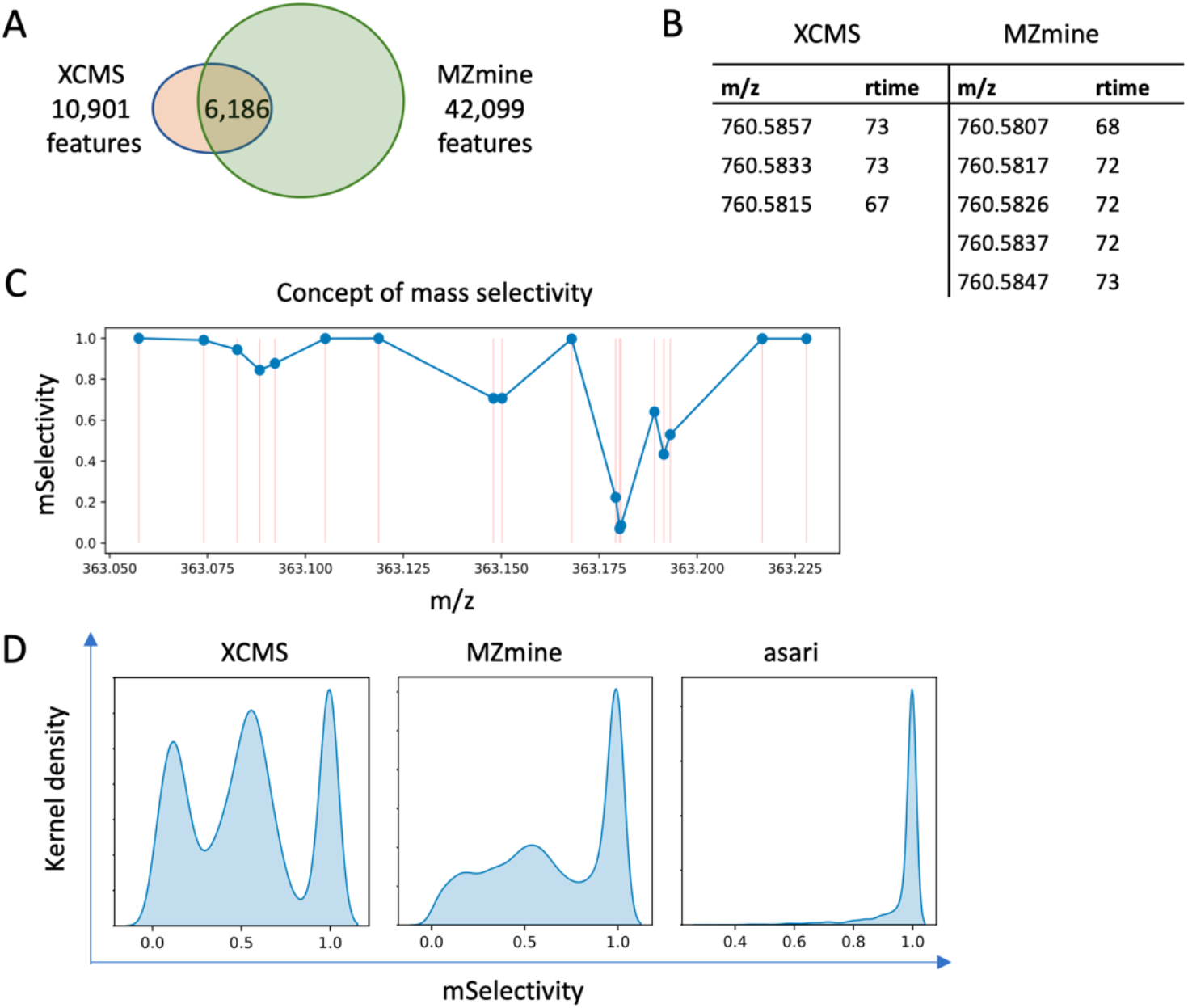
Provenance issues of feature correspondence in LC-MS metabolomics data processing. A) On a dataset of 184 repeated samples analyzed on an Orbitrap ID-X mass spectrometer (HZV029q), XCMS generated 10,901 features and MZmine generated 42,099 features (see Methods for versions and parameters). Between the two results, 6,186 features are uniquely matched. Additional comparisons are reported in **Tables S1** and **S2**. B) Many mismatched features are due to failure to resolve reciprocal best match. This example shows all 3 features in XCMS match to all 5 features in MZmine. Retention time (rtime) is in seconds. C) Illustration of mSelectivity as a function how distinct a m/z value is, regarding to its neighboring features and mass resolution. Each dot represents a m/z feature, and its mSelectivity value (Y axis) depends on the horizontal distance to neighbor features. The error in matching m/z values is modeled as a Gaussian distribution dependent on mass resolution, and mSelectivity is low when a feature has neighbors with close m/z values. D) Distribution of mSelectivity in the features produced by three processing tools. Feature m/z values are rounded to 3rd decimal place, so that minor variations are ignored and split peaks are not taken into account. Rounding has no impact on asari, because asari m/z values are linked to mass tracks.

Different from earlier studies, almost all XCMS feature here can find a counterpart in the MZmine result, within 5 ppm of m/z and 6 seconds of retention time. This is because HZV029q has better data quality than older studies, and it’s significantly larger. If a feature is missed in 1 sample, it is unlikely for the processing software to miss it in the other 183 samples. The problem here is that many features are not uniquely matched. Here, a unique match is defined as a pair of features that are reciprocally best matches in the allowed m/z and retention time windows. An ambiguous match is shown in **Figure 1B**, where 3 XCMS features are indistinguishable from 5 MZmine features. The possible reasons for this type of mismatch include a) split mass peaks on the same feature, b) failure in m/z alignment cross samples, and c) that multiple features do exist because of close chemical properties. The last scenario does occur but not in high frequency. All three scenarios violate the mass resolution of 5 ppm set for this experiment, i.e., 760.5807 and 760.5817 cannot be distinguished on the mass spectrometer. If the processing software does not resolve the issue, i.e., reporting a unique numerical value for the same mass, errors propagate into downstream annotation and statistical analysis. The confusion in comparison between tools reflects their internal confusion in feature correspondence, more specifically, mass alignment.

Using a similar concept to selectivity in mass spectrometry, we define a “mSelectivity” function to calculate how well a m/z feature is distinguished from others under a given mass resolution (see **Methods**). The concept of mSelectivity is illustrated in **Figure 1C**, where a m/z feature has a low mSelectivity score when it has neighbors of very close m/z values. We computed mSelectivity scores for the features from XCMS and MZmine, and their distributions are shown in **Figure 1D**. These mSelectivity scores were computed after their m/z values were rounded to the 3^rd^ decimal place and collapsed into unique lists, to forgive rounding errors and exclude possible isomers (i.e., compounds of the same mass). **Figure 1D** indicates that a very large number of features in XCMS and MZmine have poor mSelectivity that is not consistent with the resolution of instruments. The ideal result is that all features have mSelectivity close to 1, which is perfect compliance of mass resolution. That is how we implemented mSelectivity requirement in our new software asari (right panel in **Figure 1D**).

The distribution of mSelectivity is a straightforward summary of how well software tools distinguish m/z values in the metabolomics feature tables. The poor characteristics of XCMS and MZmine in **Figure 1D** reflect artifacts in feature correspondence, because the mass peaks detected in a single sample prior to correspondence do not have as severe a problem. Retention time in chromatography may help distinguish compounds of similar m/z values, but it does not fix issues in m/z alignment in the software, and the sheer number of problematic m/z values causes significant problems in reproducibility.

In practice, an ad hoc step could be performed to merge these close features. XCMS has a step of merging neighbor peaks that are too close, which is among the reasons of fewer features reported here. But **Figure 1D** shows clearly that it does not solve the real problem of mass selectivity. Furthermore, “manual” merging is a subjective approach that is usually in the territory of expert users. It causes issues in the quantification of these features implicitly because merging alters processing history and it does not affect all peaks equally. Most critically, manual merging does not help the reproducibility of software.

### Mass alignment should not be conditioned on elution peak detection in high resolution metabolomics

The design of XCMS and MZmine is similar: extracted ion chromatograms (EICs or XICs) are built on regions of interest (ROIs) in each sample; elution peaks are identified on each EIC; elution peaks are aligned cross samples to become features, i.e., feature correspondence (**Figure 2A**, left). Here, m/z alignment occurs after detection of elution peaks in each sample. In this approach, two peaks in a sample can start with the same m/z value but end up with different m/z values after correspondence. We note that the construction of chromatograms is straight forward in high resolution data, but detection of elution peaks is very error prone (discussed in a later section). The problem may not be pronounced in low-resolution data with few samples, but is amplified exponentially by a large number of peaks and a large number of samples. Without tracking EICs explicitly, it is a poor design to perform m/z alignment after elution peak detection.

**Figure 2.**
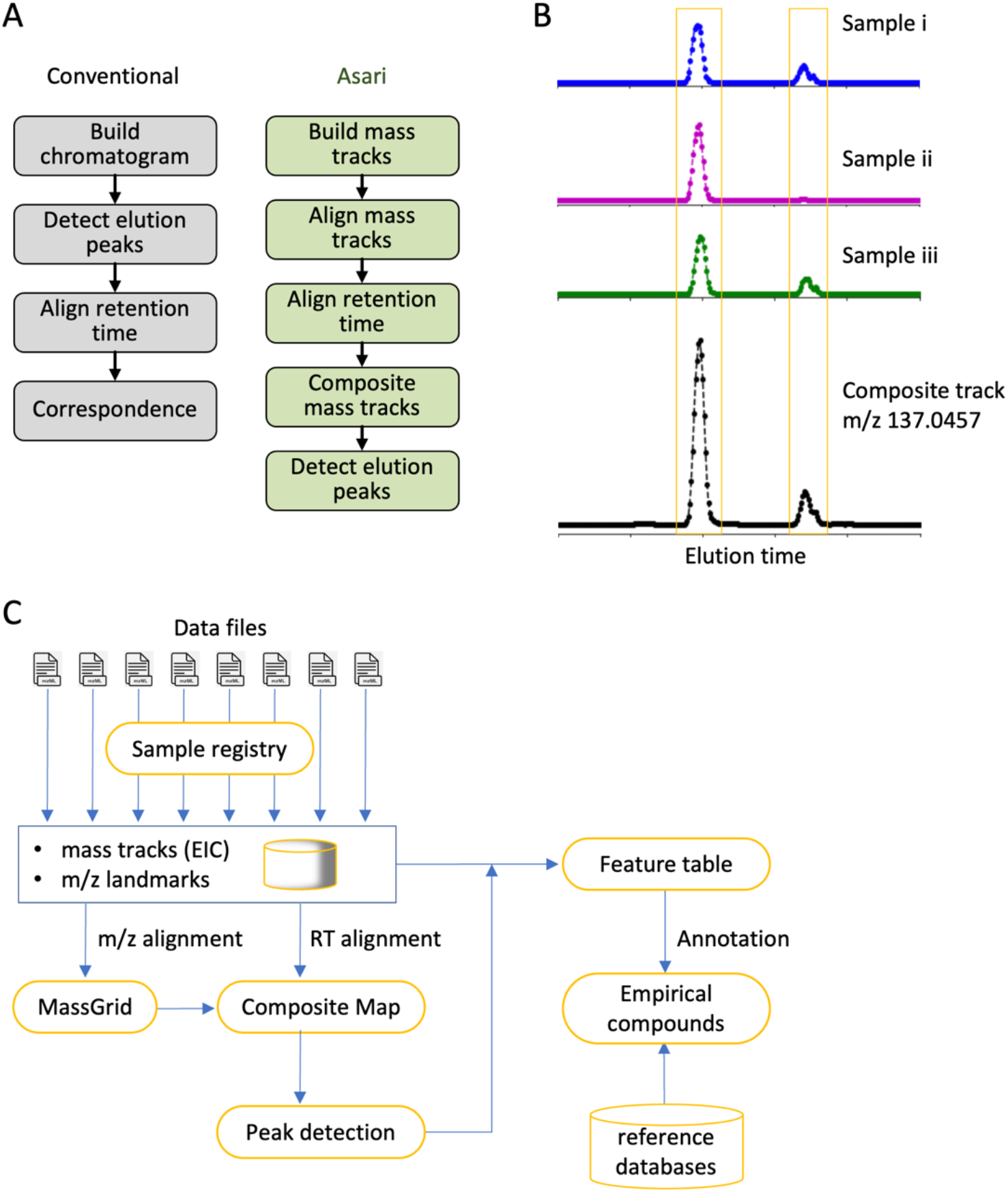
New software design of asari anchors on composite mass tracks. A)Overview of design differences between conventional software tools and asari. B)The “composite mass track” is a representation of data from all samples, by adding up the signals in corresponding mass tracks after retention time (RT) alignment. Shown are real data in MT02 dataset. C) Asari takes centroid mzML files as input, and build chromatograms for each as mass tracks. To prioritize modern mass resolution, m/z alignment is performed first to form a MassGrid, aided by isotopic landmarks. Aligned mass tracks across samples are corrected for RT, using LOWESS regression on a subset of high-quality elution peaks, then aggregated into composite mass tracks (B). All composite mass tracks are stored in the “Composite Map”. Elution peak detection is performed on the composite mass tracks, and feature table is generated by looking up the corresponding peak areas in each individual sample. Annotation groups degenerate features into empirical compounds, and reference databases are used to match the m/z values in empirical compounds.

We implemented a concept of “mass track” to address this issue. A mass track is defined as a series of LC-MS data points of the same consensus m/z value and spanning the full retention time (**Figure S1**). A mass track may or may not have detectable elution peaks, and zeros are filled for scans without an intensity value. Therefore, a mass track is an EIC that spans the full retention time, and locks the origin of peaks in computational processes. Mass tracks are aligned cross samples before elution peak detection in asari (**Figure 2A**, right). This takes advantage of the superb mass resolution on today’s instruments and avoids errors stemmed from elution peak detection. Mass tracks lock in unique m/z values per sample, therefore greatly reducing the complexity of m/z alignment, compared to the alignment of individual elution peaks. We can already use 13C/12C isotopic patterns and landmark tracks of high mSelectivity to aid the m/z alignment.

### Mass tracks and composite mass tracks anchor LC-MS alignments in asari

In biological studies, a feature is expected to have variations and it is common that it is under detection limit in many samples. Even if no elution peak is detectable, mass tracks still often exist in those samples, and the information is useful towards the correct m/z alignment. On the other hand, feature detection should utilize the recurrent pattern of the same peak in many samples. In asari, aligned mass tracks are aggregated into a “composite mass track” after correction of retention time (**Figure 2B**). With this approach, elution peak detection is no longer required on individual samples. It can be done on the composite mass track, then the peak area is looked up in each individual sample and reported as feature intensity values (**Figure 2B**).

This greatly simplifies feature correspondence, and has a significant performance gain, by not repeating the computational cost of peak detection on all individual samples. Because the composite mass track has higher signals than any individual sample, the quality of peak detection is often improved. In practical terms, detecting weak peaks will be enhanced by combining signals from multiple samples; irregularity such as chromatographic gap and bad peak shape will be ameliorated (example in **Figure S2**).

The overall design of asari is illustrated in **Figure 2C**. Aligned mass tracks of all samples form a “MassGrid”, which provides a foundation of trackable and high-quality feature correspondence. Retention time (RT) correction between samples, also called retention time alignment, is carried out by LOWESS regression, using a small subset of high-quality elution peaks. These reference peaks are easy to select from previously aligned mass tracks, preferably as the only peak on a mass track. The regression result is translated to an RT remapping function for each sample.

This remapping is bidirectional: RT on the composite mass track is mapped back to each sample correctly regardless how much error is in the regression model. The error only affects the peak range on the composite mass tracks, which is mostly inconsequential. RT alignment has no impact on m/z alignment.

Previous tools usually require the presence in multiple samples to call a feature, because of their high error rates. If a feature is only present in one or few samples, correspondence becomes problematic in these tools. However, the presence of a single good elution peak is evidence that the analytical method is valid for that particular metabolite. It is important to know that weak or no peaks in other samples is reflecting biology not inadequate chemical analysis. Since asari considers all mass tracks in the composite map, confidence can be established at high level of sensitivity. Even a peak is only present in a single sample, it will be detected and reported in asari, which is important to applications such as personalized medicine and exposomics.

### Composite map enables easy and interactive inspection of data

The design of asari significantly improves provenance through data processing. To track a problem in software, computational objects in each step have to be explicitly linked while each step shall be verified separately. Currently, it is cumbersome to verify peaks from XCMS. It usually requires scripting to plot each ROI associated with a feature, which is not real backtracking because an ROI is not the same as EIC and has more data points. On the other hand, the computational complexity increases quickly if all intermediate steps are recorded (part of the performance penalty in MZmine). By linking features and raw data through composite map, asari produces an efficient solution of automated testing, debugging and verification. A significant change is that asari composite map is representative of all samples – the burden of peak evaluation and visualization is largely removed from repeats on individual samples. Therefore, this enables a data dashboard for users to navigate and inspect data, regardless of the size of samples.

An example asari dashboard is shown in **Figure 3A**. This is an interactive tool that users can launch into a web browser after each dataset is processed. The top part of the dashboard is a set of tabs for data summary and quality metrics. Users can navigate through the feature browser to inspect EICs by clicking, hovering, panning and zooming functions. A separate mass track viewer shows all detected peaks on a mass track, with an example screen shot shown in **Figure 3B**. When users zoom in different regions of the mass track, the peaks can be inspected closely (**Figure 3C, D**). The ability to inspect data and feature quality visually is important for the reproducibility of science, and should be done before resources are committed to continued work. The asari dashboard is not only useful to end users, but also to developers and data scientists for testing and interacting with the software.

**Figure 3.**
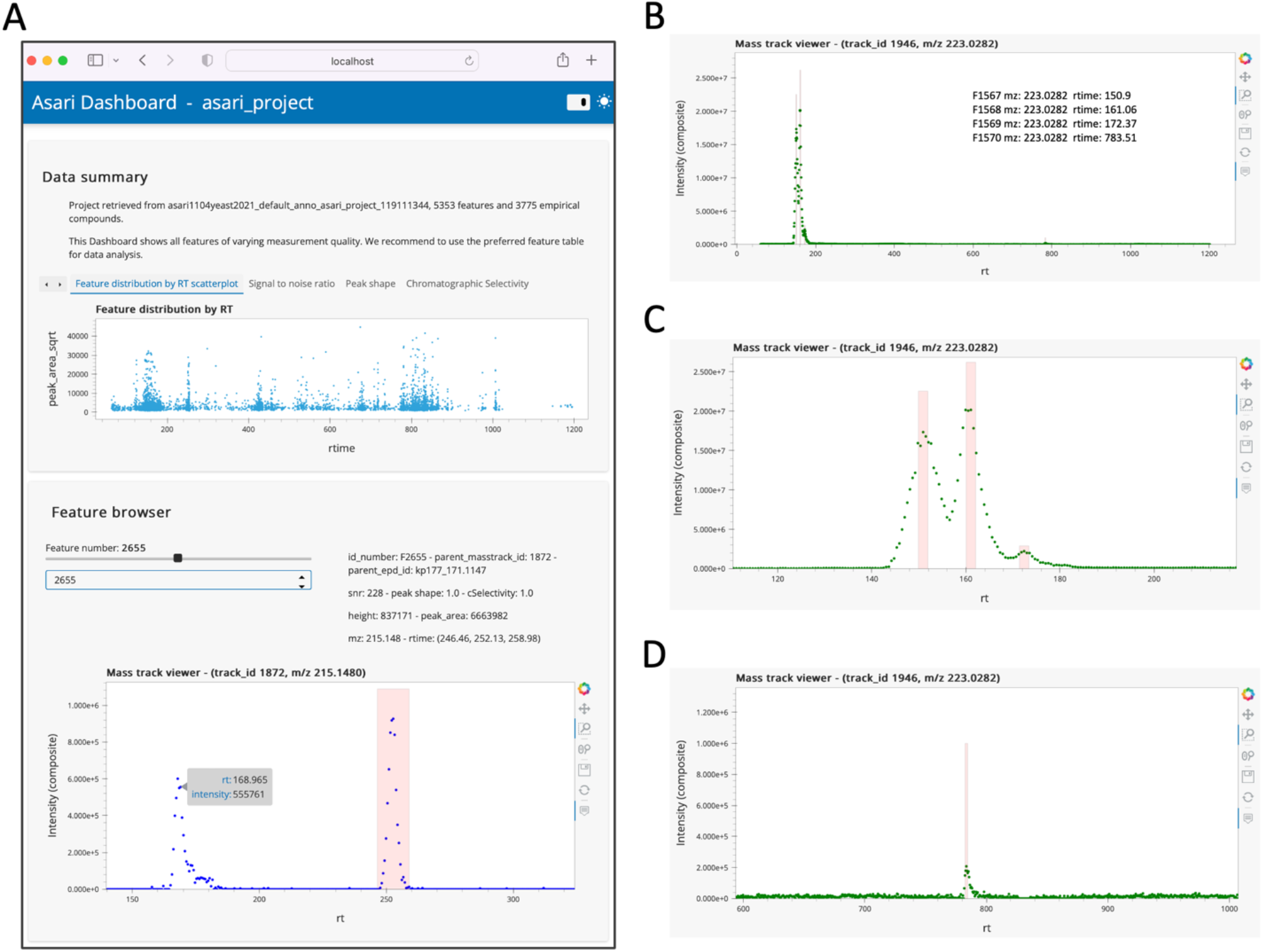
Composite map enables easy and interactive inspection of data in a web browser. Users can click, zoom, pan and hover on the interactive figures. Raw data points are plotted without smoothing. The dashboard can be launched by asari viz subcommand. A) Screen shot of asari Dashboard, with the Feature browser. Users can click through all features or find by feature ID. B) A mass track with four peaks, and zoom-ins are shown in C) and D). Mass track viewer supports m/z search.

### Asari provides verifiable elution peak detection

The new design of asari leads to the improvement of multiple key aspects. We start with benchmarking feature detection, before discussing the underlying mechanistic of elution peak detection and quality metrics. Of note, the approach here on feature detection does not concern false positives, which are discussed in the next section. The benchmark datasets have to be impartial to the tools being tested.

We used the manufacturer’s software to process, select and visually verify 402 features in the HZV029 data. The detection of these features by each tool is reported in **Figure 4A**. In its default setting, asari detected 386 of these features, the second highest next to MS-DIAL. Of the 16 features asari disagreed with the manual list, 7 were on low quality EICs discarded by asari, and 9 were peaks of poor shape and borderline number of data points (**Figure S3**). We note that the higher number of detected features by MS-DIAL is attributed to its low stringency by reporting 29,924 features, several times more than others. All the 8 peaks missed by MS-DIAL, however, have good peak shape and were detected by asari (**Figure S4**).

**Figure 4.**
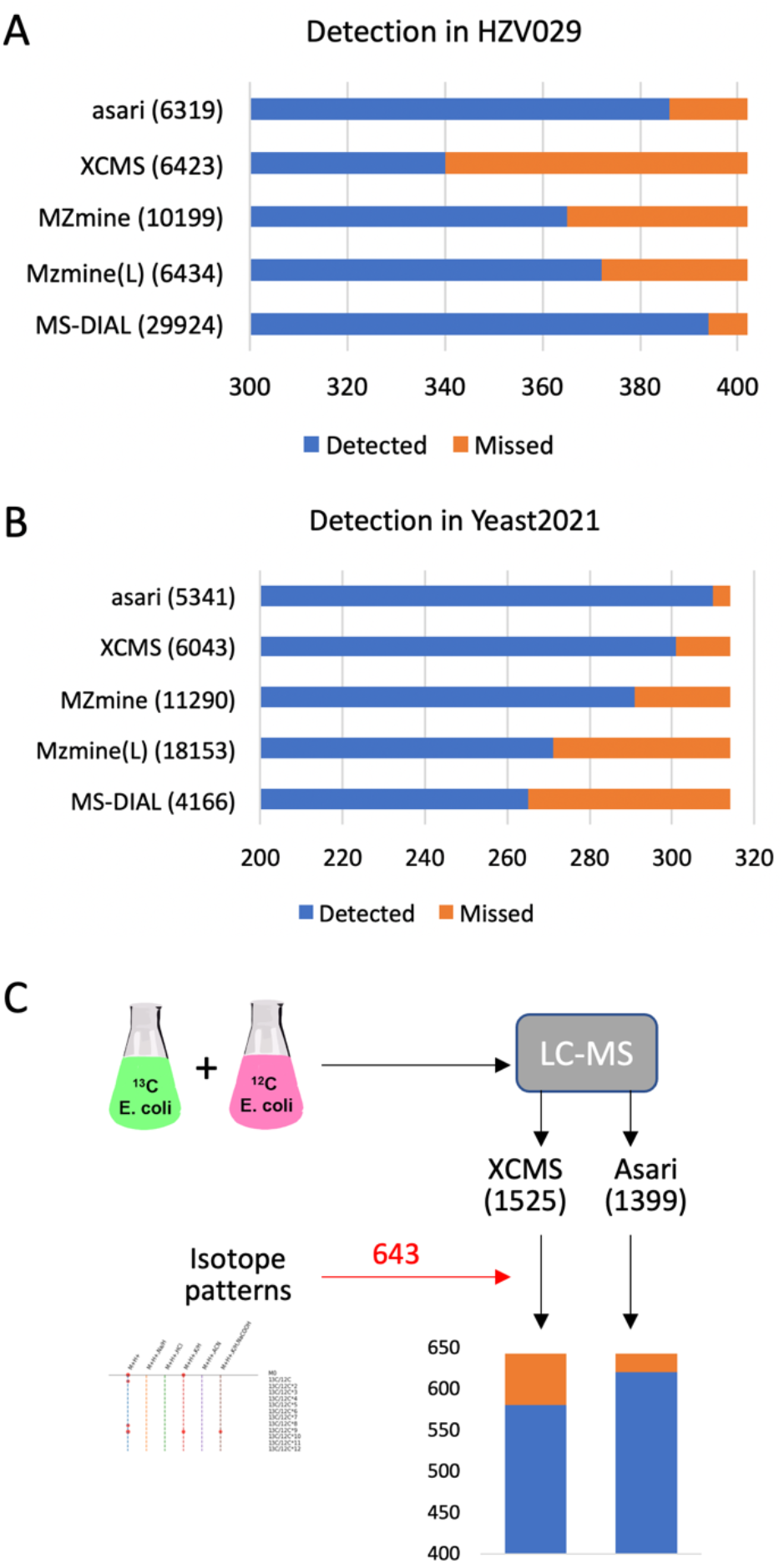
Evaluation of asari feature detection. A) Detection in HZV029q dataset, based on three randomly selected samples. The manually verified 402 features were selected from results by Thermo Scientific software. B) Detection of manually certified features in three samples in Yeast2021 dataset (Chen et al, 2021). MZmine version 3.3.0 (wavelets) produced identical results as version 2.53 in both datasets (A and B). C) Detection in the credentialed E. coli data. XCMS and asari used similar parameters (peak height > 1E5). The two feature tables were annotated separately using khipu (Li et al, 2023) to identify isotopic patterns. All [M+H]+ features with correct isotopic patterns were combined into the 643 “certified” features. Missed feature numbers are 62 for XCMS and 22 for asari (further examined in **Figure S6**).

A second benchmark dataset is Yeast2021, in which Chen et al (2021) manually confirmed 314 identified features. The detection result by each tool is shown in **Figure 4B**. Asari returned the best performance of detecting 310 of the 314 features. The 4 features missed by asari are shown in **Figure S5**: two of them have too few data points, one under minimal requirement of peak height, and one with too high local noise.

Additional comparisons were performed between XCMS and asari. We have analyzed the human plasma reference sample NIST SRM 1950, and verified 39 metabolites previously reported in the literature (Simon-Manso et al, 2013). Both asari and XCMS successfully detected all these 39 metabolite features (see Methods). We generated a new dataset based on credentialed E. coli samples (**Figure 4C**, similar to Mahieu et al, 2014). A subset of E. coli metabolites was labeled by ^13^C isotope during the cell culture. XCMS and asari extracted 1525 and 1399 features from this dataset, respectively. After matching isotopic patterns between labelled and unlabeled samples, 643 features in total showed correct isotopic patterns as protonated ions. Among them, XCMS detected 581 and asari 621. Among the 22 features missed by asari, two features were out of the 5 ppm m/z range; the other 20 are plotted in **Figure S6**. Three of them would pass as real peaks with more aggressive smoothing, and the remaining 17 features do not meet quality requirements in asari. Taken together, these results indicate that the peak detection performance of asari compares favorably against others, and the behavior of asari is understandable and predictable.

### The goal of metabolomics peak detection is not more features but fewer errors

Elution peak detection deserves a detailed investigation, as so many confusions are related to peak detection as well as the coverage of metabolomics. As seen in **Figure 1** and **Tables S1** and **S2**, the numbers of features are quite different from different tools, even though XCMS and MZmine have the same wavelet algorithm (Tautenhahn et al, 2008). Therefore, the differences are not entirely caused by peak detection algorithms. We discussed above the impacts from mass alignment and chromatogram construction. When different peak detection algorithms are used within MZmine, using the same EICs, they still returned very different numbers. It’s a complex problem involving many components in software implementations. Just as in other – omics data, metabolomics features should be considered in a statistical context.

To give a visual guide to the problems, we illustrate a few typical EICs in **Figure 5A**, In the “good” cases, almost any peak detection algorithm will work. In other cases, the consideration of noise level is critical. In high-noise EICs, one can get more peaks by lowering stringency of parameters. An extreme example is shown in **Figure S7A**, where the number of peaks inflates quickly by varying parameters. We have to caution that sensitivity cannot come at the cost of low data quality. At low noise level, small peaks can be valid (**Figure 5A**, upper right). A peak detection algorithm should ensure they are detected even at the presence of multiple big peaks.

**Figure 5.**
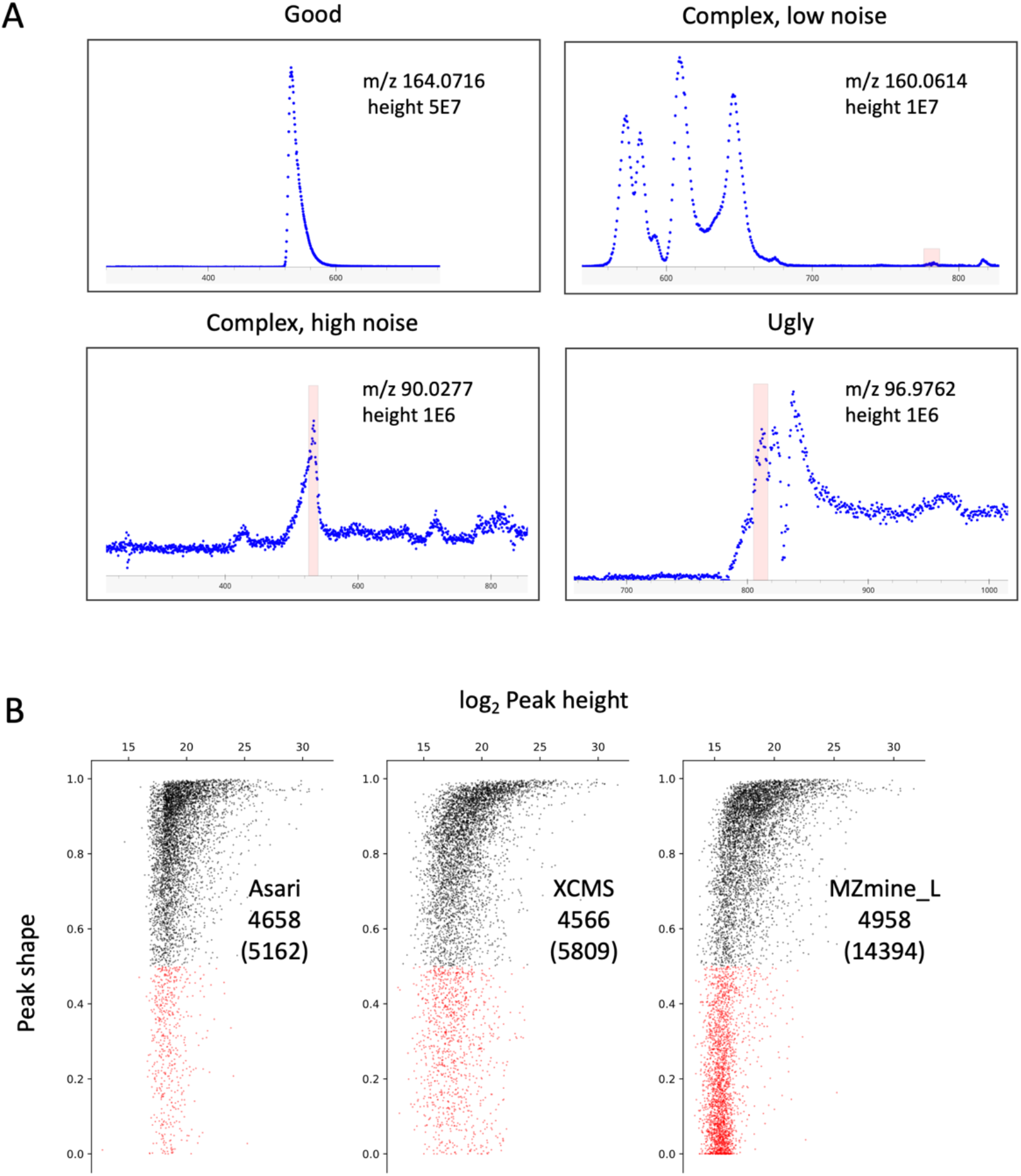
The number of peaks has no significance beyond a core set of high-quality peaks. A) Examples of different types of EICs. Levels of maximum height, baseline and noise are used to inform algorithmic decisions involved in peak detection. B) Re-computed peak shapes for features reported by different tools on the Yeast2021 dataset (see **Table S2**), plotted as a function of peak height. Each dot represents a LC-MS feature, black for Gaussian fitness score > 0.5. C) MS-DIAL is not included in this analysis because we did not find a straight forward method to export peak boundaries after alignment. For other tools, all features from each were mapped to a set of mass tracks generated by asari using low thresholds (minimal peak height 5000, peak shape 0.1, SNR 1.1). Almost all m/z values in the tools were matched to these mass tracks. For each feature, the retention time was padded 3 seconds on each side of peak boundary, to accommodate processing variations. D) The redundant features (within 5 ppm and 6 seconds) were first grouped in XCMS and MZmine results. Thus, the total feature numbers in this analysis are 5809 for XCMS, 10876 for MZmine (wavelets), and 14394 for MZmine (local minimum). The MZmine (wavelets) result has 6841 good peaks and 4035 bad peaks. They are not plotted here because too many low intensity peaks show as 0s in this approach.

The default peak detection algorithm in asari first estimates the baseline and noise level on a mass track. The statistics drive the decisions on baseline subtraction, detrending or smoothing. Detrending is to regress out shifting chromatographic background, used sparely in necessary tracks (e.g. **Figure S7B**). Noisy mass tracks are smoothed using simple moving average. Aggressive smoothing can lead to superfluous artifacts (the reason that raw data points have to be used for inspection) and is not recommended for general use. A mass track is then separated into segments of valid signals, and peak detection is performed on each segment. We use a local maximum search algorithm, with peak height and prominence requirements dynamically determined on the segment statistics. Prominence is the vertical distance of a peak top to its adjacent local minima, therefore important to control for fluctuating data points. The detected peaks are assessed by a set of quality metrics and retained only if the preset thresholds are met. In larger studies, asari also benefits greatly from the cumulative peak patterns in composite mass tracks.

To understand the vast differences between tools (**Tables S1, S2**), we need to ask how many good peaks are found by them. An intuitive measure of peak quality is “peak shape”, here defined as the goodness of fitting to a Gaussian curve (other curves have been used but the impact on fitness scores is negligible). High quality peaks should have peak shapes close to 1. We recomputed the peak shapes from the result of each tool on the Yeast2021 dataset (longer chromatography, small size of only three samples minimizing impact from mass alignment), and plot them as a function of peak height in **Figure 5B**. Asari by default does not return peaks of bad shapes; the 504 red features were due to the padded retention time (to make features comparable across tools) in the recomputing of peak shape. XCMS has 1243 features in red; MZmine (wavelets) has 4035 in red (not shown); and MZmine (local minimum) has 9436 in red. This indicates that the number of good features is stable. The large numbers of “extra” features reported by different tools are low-quality peaks, which are not reproducible and should not be used without additional evidence. Indeed, the overlap features between asari and others are stable at around 3000 (**Tables S2**). **Figure 5B** has close to 5000 in black, because not all features are uniquely matched, due to the correspondence problem discussed earlier. Summarizing results in **Figures 4** and **5**, larger numbers of reported features do not improve the real performance on benchmark peaks or good peaks.

### A set of metrics fully describe data quality

The discussion above shows the importance to safeguard the processing result by quality metrics. High numbers of bad peaks are not acceptable in real-world applications. They impact projects very negatively. In biomedical studies, features of questionable quality cannot be used for decision making. Computational processing cannot overstate the quality of analytical chemistry.

SNR (signal-to-noise ratio) is often defined differently among tools. In asari, the noise for an elution peak is the average signal intensity of neighboring non-peak data points, 100 to each side; SNR is the ratio between peak height and noise (**Figure 6A**). The mSelectivity metric (**Figure 1C**) is baked into asari algorithms. Similar for chromatography, cSelectivity is defined as how distinct chromatograhic elution peaks are (**Figure 6B**). Together with peak shape, users can rely on them to determine the peak quality on a daily basis. Global visualization of these three quality metrics in Yeast2021 data, in relation to peak size, is shown in (**Figure 6C**). This figure demonstrates the desired quality of good peak shape, high SNR and high cSelectivity on asari features. The same plot can be used to weed out poor data quality. The SNR distribution is an informative way to assess how much useful information is in a metabolomics dataset (**Figure 6D-F**). These quality metrics are exported with the feature tables in asari. With their assistance, users own the decisions of filtering data, and full confidence in the feature quality.

**Figure 6.**
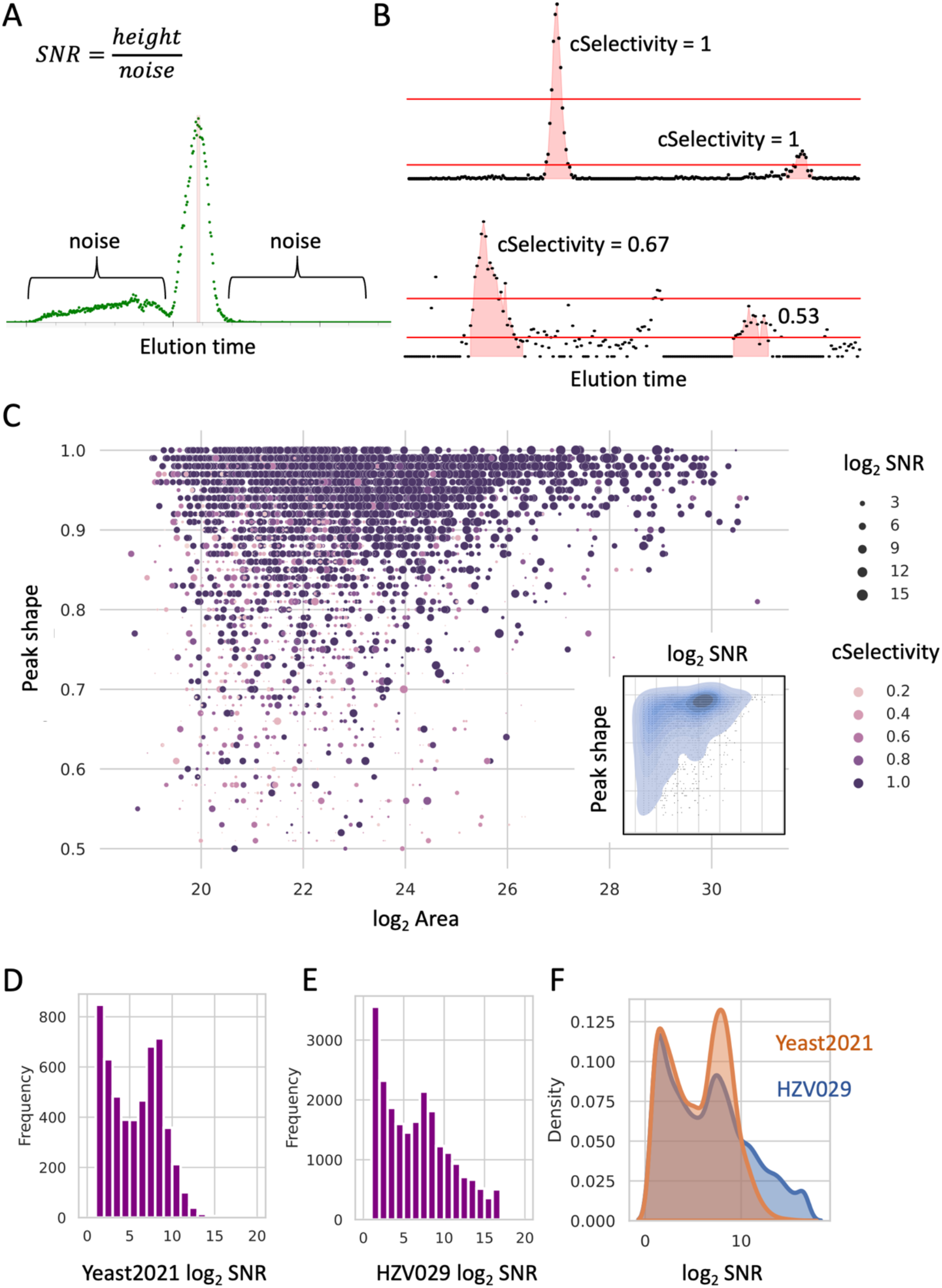
Quality metrics in asari and applications to data review. A) Signal-to-noise ratio (SNR) in asari is defined by peak height divided by noise level. In a mass track (extracted ion chromatogram), all data points within the peak range are considered part of the peak. All other data points are considered as noise. The noise level is taken as average intensity of up to 100 nonpeak data points on each side of the peak. B) Chromatographic peak selectivity (cSelectivity). After filtering the data by 1/2 of this peak height, the cSelectivity of this peak is defined by the fraction of the data points belong to any peak. cSelectivity is 1 when the chromatogram has no noise above the half height of any peak. C) Overview of all features in the Yeast2021 dataset. Peak area (log2) and peak shape are x and y-axes; SNR and cSelectivity are coded by size and color. The inset shows the kernel density of SNR and peak shape. D-E) Histograms of SNR distribution in the Yeast2021 and HZV029 datasets. F) Kernel density of SNR distribution in the Yeast2021 and HZV029 datasets. More features with high SNR indicate better quality in feature detection.

### Feature quantification is evaluated favorably in asari

The quantification of each peak is usually based on the peak area, which is represented in asari by summing the intensity of each data point in a peak. The peak areas by XCMS and asari are generally agreeable (**Figure 7A-B**). To further investigate the performance in quantification, we designed an experiment where human plasma and vegetable juice were mixed by varying ratios (BM21 dataset, **Figure 7C**). Therefore, a majority of features are expected to have their peak areas correlated with the mixing ratios. Overall, 5581 features were matched between XCMS and asari in the BM21 dataset. Their Pearson correlation coefficients to the mix ratios were computed, and the distributions are shown in **Figure 7D**. Asari has more features with correlation coefficient > 0.9, indicating that the quantification in asari is more useful than that of XCMS.

**Figure 7.**
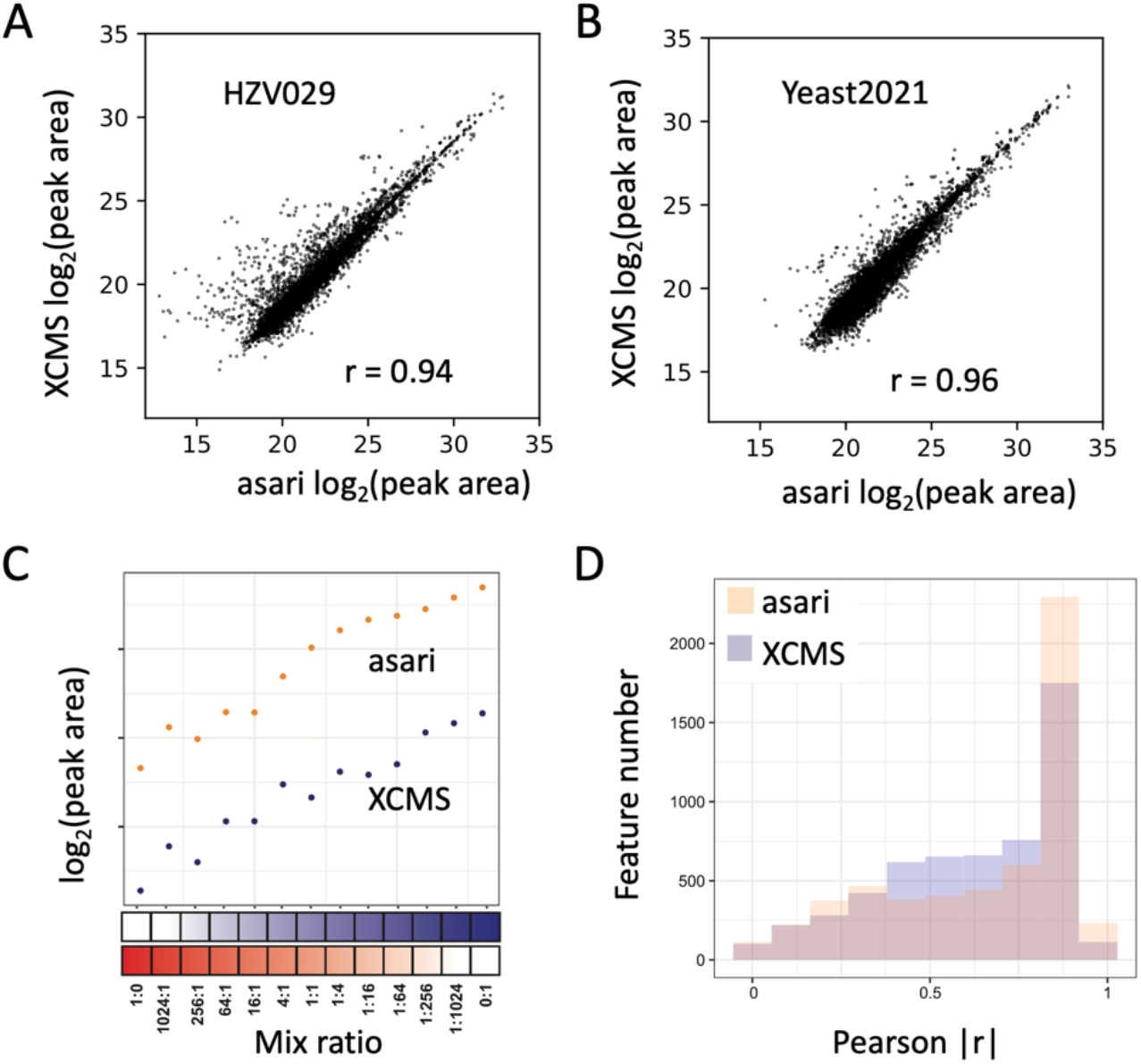
Evaluation of feature quantification. A-B) Scatter plot of the log2 peak areas of common features between asari and XCMS, on three samples in HZV029 dataset (A) and Yeast2021 dataset (B). The r values are based on Pearson correlation. C)Design of the BM21 dataset, by varying mix ratios between human plasma and vegetable juice. A well quantified metabolite is expected to show good correlation between the mixing ratios and the reported peak areas, as exemplified by the feature on top (m/z 189.1232, 159 seconds). Asari calculates peak area differently from XCMS, resulting in higher values in Orbitrap data. D)Overall quantification results in the BM21 dataset, shown as feature numbers binned by Pearson correlation coefficients between peak areas and sample mixing ratios.

### Asari delivers a new generation of computational performance

Computational efficiency is fundamentally important to –omics data. For MZmine and MS-DIAL, processing 100 samples often becomes challenging on a desktop computer. XCMS is considered more performant and is the default choice of cloud computing (Tautenhahn et al, 2012, Delabriere et al, 2021, Pang et al, 2021). The computational efficiency of asari is benchmarked against XCMS here using multiple datasets of different sizes and platforms. Small studies can be processed by asari under a minute. Some studies of 100∼200 samples take less than 10 minutes by asari using a single CPU core. Therefore, asari provides significant improvement of CPU time over XCMS by 1∼2 orders of magnitude (**Figure 8A**).

**Figure 8.**
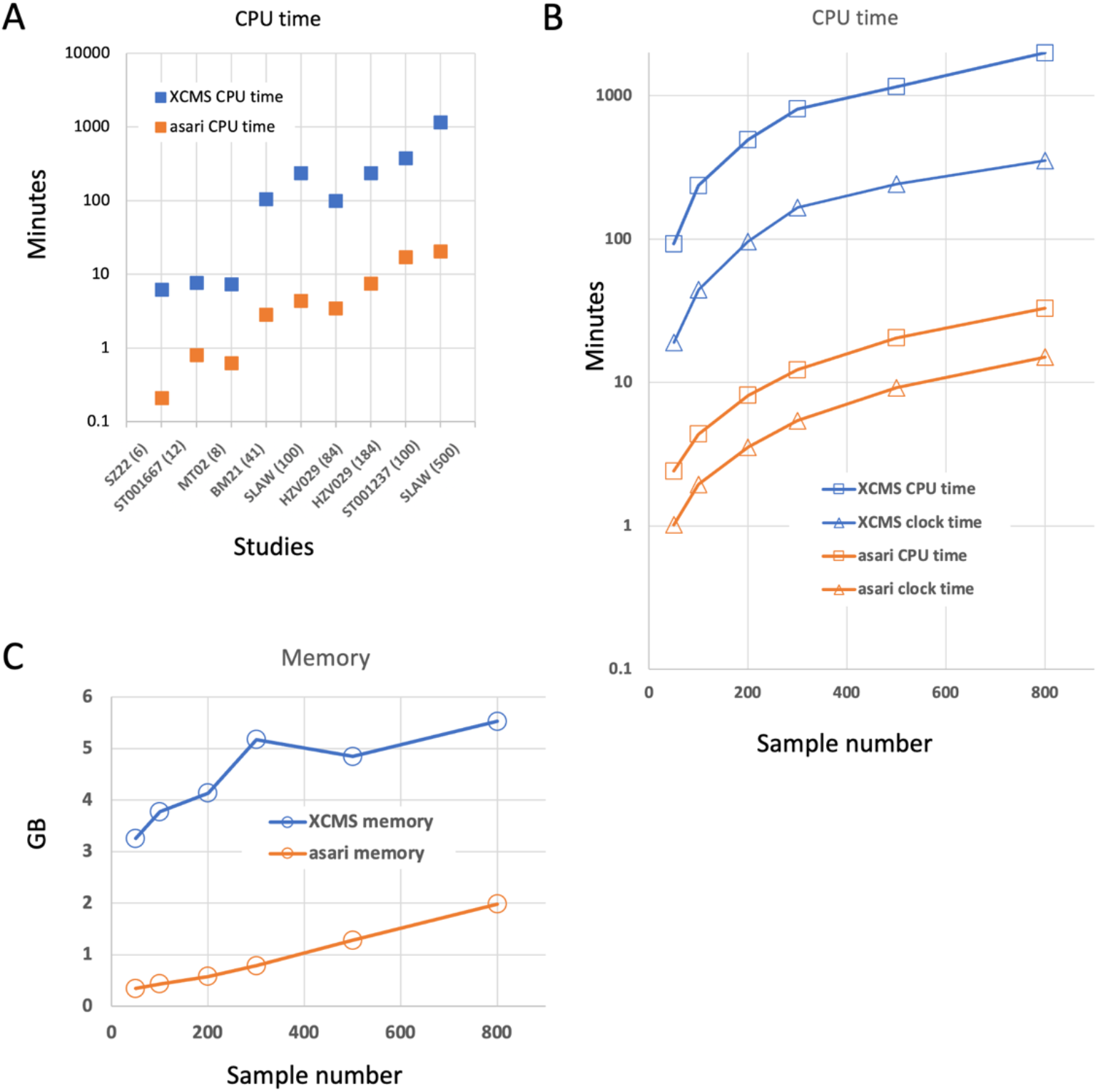
Evaluation of computational performance. A) Computational performance in user CPU time (equivalent to single core) by asari and XCMS on different datasets (sample numbers show in parentheses on X-axis). B-C) CPU time and wall clock time (B) and memory (C) used by asari and XCMS on the SLAW dataset using varying number of samples. Y-axis is in log10 scale for CPU time.

To test the scalability, we subset the SLAW data (Delabriere et al, 2021) using varying sample numbers. The CPU time and memory use is mostly a linear function of sample numbers (**Figure 8B, C**). The results indicate that the performance gap between XCMS and asari widens for larger studies. XCMS can also become more complicated if it goes beyond simple workflows or large studies are processed (Delabriere et al, 2021). The full SLAW dataset of > 2,000 samples was processed in the previous study by XCMS on a cluster node of 15 CPU cores in 7∼12 hours. Now it takes asari ∼1 hour on a regular laptop computer. These results indicate that asari delivers a new generation of computational performance. This enables expeditious metabolomics data processing on cheap hardware, and makes very large studies feasible.

### Reusable data structures and code, a bridge to data science

The asari software is built on a set of transparent data structures (**Figure 2C**). Mass tracks are extension to the concept of extracted ion chromatograms, to simplify m/z alignment and navigation. A MassGrid records the alignment of mass tracks across samples. A feature is defined at experiment level, and elution peaks are defined at sample level. Mass tracks cross samples are superimposed and summed up, to become composite mass tracks. The composite mass tracks are representative of all samples. A metabolite may have multiple degenerate features due to isotopes, adducts, neutral loss and fragments, which are grouped into an “empirical compound” via another package khipu (Li and Zheng, 2023). An empirical compound is a computational unit for a tentative metabolite, since the experimental measurement may not separate compounds of identical mass (isomers). Asari explicitly links mass tracks, peaks, features and empirical compounds, so that each processing step can be traced and verified. Examples of data structures are provided in **Methods** and in our code repositories. These data structures are JSON compatible and exposed, so that advanced users and developers can reuse them easily.

We designed asari with the goals of reusable code, easy deployment and maintenance, being cloud friendly and scalable performance. The software is open source on GitHub, available via standard Python package management tools, which can be integrated seamlessly with cloud deployment. Docker images and build recipe are available. Subcommands are designed to perform common tasks, such as batch processing, analyzing experimental parameters, targeted extraction, and visualization via the dashboard. The software modules can be ported for other tools or scripting, as demonstrated by Jupyter notebooks in the code repositories.

As metabolomics is here to serve biomedicine, metabolomics data processing tools have to be used by general data scientists, not limited to chemists. Asari marks a clear transition: previous tools are burdened with complicated parameters while asari requires almost no tunable parameters. To meet the diverse demands of advancing science, the software has to be interoperable, and interface with the rich tools in general data science. The above features of asari are designed to bridge the gap to data science, and fit easily into automated pipelines.

## Discussion

Reproducibility of data processing has been a roadblock of metabolomics. In this study, we attribute the inconsistency in previous software tools to poor mass alignment and lack of quality metrics. The widely used wavelet algorithm shows no advantage in this study, but costs unnecessary complexity. We have developed the asari software from ground up, which compares favorably to others in feature detection. Arguably, the detection performance of the previous tools may be improved by further parameter tuning. But this study highlights that the number of high-quality features is stable, and increasing feature numbers brings undesired false positives.

Guarded by a more statistical approach, asari does not require users to supply any tunable parameter than mass resolution, where the default value of 5 ppm rarely requires changing. Previous software tools often deteriorate in feature correspondence in larger studies. On the contrary, the processing quality in asari increases with larger studies, because it utilizes recurrent patterns in feature detection. Asari has been mostly tested on the Orbitrap platforms. Continued development and community involvement will be important to cover the diverse platforms and methods in metabolomics. For example, additional methods of retention time alignment can take advantage of spike-in standards. Lipidomics and xenobiotics can benefit from additional specific modules. There have been debates on the use of absolute amu or relative ppm for mass resolution in data processing. Asari uses ppm currently. We note that the former is 0 order and the latter 1^st^ order of a polynomial model, but a 2^nd^ order polynomial model may be required to cover the full m/z range.

In summary, the development of asari has significantly contributed to the reproducible data in metabolomics, by a full set of linked and transparent data structures in all processing steps. This allows developers to trace, debug and optimize the process into the future. The end users can navigate and verify features by interactive visualization of extracted ion chromatograms in asari dashboard. Asari has delivered a new generation of computational performance, which is necessary for the future growth of metabolomics.

## Data availability

The BM21 and HZV029 datasets are available at Metabolomics Workbench (https://www.metabolomicsworkbench.org), by IDs ST002233 and ST002454, respectively. The datasets MT02 and SZ22, and the list of verified NIST SRM 1950 features, are available at https://github.com/shuzhao-li/data. The large SLAW dataset (Delabriere et al, 2021) was retrieved from MassIVE (https://massive.ucsd.edu) by study ID MSV000086486. The Yeast201 data (Chen et al, 2021) were retrieved from MassIVE by study ID MSV000087434. The other public datasets used in this work are under Study IDs ST001667 and ST001237 on Metabolomics Workbench.

## Code availability

The asari source code is available at GitHub, https://github.com/shuzhaoli/asari, and as a Python package via https://pypi.org/project/asari-metabolomics/. Jupyter notebooks used for data analysis in this paper are provided at GitHub, https://github.com/shuzhao-li/data.

## Acknowledgements

This work was in parted funded by NIH grants (to SL) U01 CA235493 (NCI) and R01 AI149746 (NIAID).

## Author contributions

S.L. designed the study, wrote the asari software and the manuscript. A.S. and S.L. performed data analysis and software testing. M.T. performed the experiments of HZV029, MT02 and BM21. S.Z. performed the experiment of SZ22.

