## Supplemental Information for "Trackable and scalable LC-MS metabolomics data processing using asari"

### Methods

**Software design of asari.** Asari is written in Python 3, and can be used as a standalone command line tool or imported as a package. Its library dependency includes numerical computing via numpy and scipy, data wrangling via pandas, and visualization via panel and hvplot. Pymzml is used to parse mzML format. Data structures, annotation, search and chemical calculation make use of our supporting packages metDatamodel, mass2chem and jms-metabolite-services. Implementation of new and previous algorithms was coded from ground up, where numerous details contributed to the computing speed, e.g., discrete mathematics is preferred over continuous curves, and intermediary indexing and caches are employed. Processed mass tracks are cached on disk to reduce memory footprint. Mass tracks are explicitly linked with features and peaks, and the information is exported as JSON in asari output. The quality metric mSelectivity is used internally. Other quality metrics, peak shape, SNR and cSelectivity, are part of exported feature tables. Grouping ions into empirical compounds is performed using khipu (Li and Zheng, 2023). The annotation and search functions are generic to accommodate reference databases, and the default is HMDB (Wishart 2020).

**mSelectivity**. The probability of confusing two m/z values *i* and *j* is modelled as

$$P_{i,j} = e^{\frac{-\Delta m/z}{\xi}}$$

, where ξ is the preset mass resolution, e.g. 5 ppm. The mSelectivity of m/z value *i* among a list of *n* values is defined as

$$\prod_{1 \le k \le n,\ \ k \ne i}^{n} (1 - P_{i,k})$$

. In practice, we only need to calculate the mSelectivity based on the two neighbors of higher m/z values and two of lower m/z values to obtain an accurate approximation.

**Metabolomics datasets**

Four new datasets were generated in this study: HZV029 (human plasma samples), MT02 (human plasma samples), SZ22 (E. coli samples with 13C isotope labeling), and BM21(serial mixture of human plasma and vegetable juice). HZV029 is used as two parts: 184 samples in HZV029q, and the other part contains a different biological sample. Four public datasets are used here: Yeast2021 (Chen et al, 2021), SLAW (E. coli, large Orbitrap QE data as described in

Delabriere et al, 2021), ST001237 (human sera, Orbitrap QE), and ST001667(mouse liver, Q-TOF data). Access of all data is described in “**Data availability**”.

**LC-MS metabolomics experiments**. The human plasma samples used in this study included a pooled deidentified QC sample in a vaccination cohort, NIST SRM 1950 (https://www-s.nist.gov/srmors/view_detail.cfm?srm=1950), and a commercial reference sample Qstd (Sterile Filtered Human Plasma (K2) EDTA, Equitech Bio, Inc. KERRVILLE, TEXAS). The BM21 experiment included a serial mixture of human plasma (Qstd) and vegetable juice, at the ratio of 1024:1, 256:1, 64:1, 16:1, 4:1, 1:1, 1:4, 1:16, 1:64, 1:256 and 1:1024. Along with the 11 serial mixture samples, 100% vegetable juice and 100% plasma were also included. All samples were analyzed in triplicates, while one replicate was used for data analysis in this study for simplicity. The dry extracts of unlabeled and $^{13}C$ labeled *E. coli* (Cambridge Isotope Laboratories, Inc.; Catalog number: MSK-CRED-DD-KIT) were reconstituted in 100 µL of ACN/$H_2O$ (1:1, v/v) then sonicated (10 mins) and centrifuged (10 mins at 13,000 rpm and 4°C) before overnight incubation at 4°C. The supernatant for each $^{12}C/^{13}C$ *E. coli* extract was collected and then prepared for LC-MS analysis. These samples were run in triplicates.

Metabolites extraction was carried out by protein precipitation technique using extraction solvent, acetonitrile:methanol (8:1, v/v) containing 0.1% formic acid and isotope labelled Trimethyl-13C3]-caffeine, [13C5]-L-glutamic acid, [15N2]-Uracil, [15N,13C5]-L-methionine, [13C6]-D-glucose and [15N]-L-tyrosine as spike-in controls. 30 ⌠l of plasma samplewas taken and 60 ⌠l of extraction solvent was added. Extraction blanks were also prepared to remove features of non-biological origins. All samples were vortexed and incubated with shaking at 1000 rpm for 10 min at 4°C followed by centrifugation at 4°C for 15 min at 15,000 rpm. The supernatant was transferred into mass spec vials and 2 µL injected into UHPLC-MS.

All samples were maintained at 4 °C in the autosampler, and analyzed using a Thermo Scientific Orbitrap ID-X Tribid Mass Spectrometer coupled to a Thermo Scientific Transcen LX-2 Duo UHPLC system, with a HESI ionization source, using positive and negative ionizations. The MS settings are: spray voltage, 3500 V; sheath gas, 45 Arb; auxiliary gas, 20 Arb; sweep gas, 1 Arb; ion transfer tube temperature, 325 °C; vaporizer temperature, 325 °C; mass range, 80-1000 Da; maximum injection time, 100 ms. The resolution was set at 120,000 in the HZV029 experiment, 60,000 in the BM21 and SZ22 experiments.

Data were acquired using hydrophilic interaction liquid chromatography (HILIC) positive and reversed phase (RP) negative polarities in full scan mode with mass resolution of 120,000 simultaneously. An AccucoreTM-150-Amide HILIC column (2.6 µm, 2.1 mm x 50 mm) and a

Hypersil GOLDTM RP column (3 µm, 2.1 mm x 50 mm) maintained at 45 ºC were used for chromatographic separation. 0.1% formic acid in water and 0.1% formic acid in acetonitrile were used as mobile phase A and B respectively for RP acquisition. 10 mM ammonium acetate in acetonitrile:water (95:5, v/v) with 0.1% acetic acid as mobile phase A and 10 mM ammonium acetate in acetonitrile:water (50:50, v/v) with 0.1% acetic acid as mobile phase B were used for HILIC method. For HILIC acquisition, following gradient was applied at a flow rate of 0.55 ml/min: 0-0.1 min: 0% B, 0.10-5.0 min: 98% B, 5.00-5.50 min: 0% B and 4.5 min for cleaning and equilibration of column. For RP column, following gradient was applied at a flow rate of 0.4 ml/min: 0-0.1 min: 0% B, 0.10-1.9 min: 60% B, 1.9-5.0 min: 98% B, 5.00-5.10 min: 0% B and 4.9 min cleaning and column equilibration. The chromatographic run time was 5 min followed by 5 min washing step after each sample.

**LC-MS metabolomics data processing**.

Asari has default parameters with mass accuracy of 5 ppm, minimum peak height 1E5. No parameter was modified unless described specifically. The results in this paper were based on version 1.10.6.

MZmine 2.53 (or version 3.3.0) processing was performed in the following order:

1. Mass detection, using Centroid mass detector.
2. ADAP Chromatogram builder, using min group size of 6 scans, Group intensity threshold 1E3, Min highest intensity 1E4, m/z tolerance 0.001 or 5 ppm.
3. Feature detection, Chromatogram deconvolution, m/z center calculation: median, using either
    a) Wavelets (ADAP) algorithm, S/N threshold 7, S/N estimator intensity window SN, min feature height 1E4, coefficient/area threshold 100, peak duration range 0.02-1, RT wavelet range 0.02-0.5. Or
    b) Local minimum search algorithm, with Chromatographic threshold 80%, Search minimum in RT range 0.02 min, Minimum relative height 5%, Minimum absolute height 1E4, Min ratio of peak top/edge 3, Peak duration range 0.02-0.5 min.
4. Alignment, Joint Aligner, using m/z tolerance 0.001 or 5 ppm, Weight for m/z 80, Retention time tolerance 0.2 min, Weight for RT 30.

XCMS (version 3.18.0) processing was performed using the following key parameters:

CentWaveParam(peakwidth = c(1, 30), ppm = 5, noise = 1000, prefilter = c(3, 1000))
MergeNeighboringPeaksParam(expandRt = 5, ppm = 1, minProp = 0.5)
groupChromPeaks: (minFraction = 0.1, bw = 3, binSize=0.001)

The R script used in the processing is posted in our asari GitHub repository under doc/.

MS-DIAL (version 4.90) processing used the following parameters:

Smoothing method: LinearWeightedMovingAverage, Smoothing level: 3, Minimum peak width: 6, Minimum peak height: 5000, Retention time tolerance: 0.05, MS1 tolerance: 0.001, Retention time factor: 0.3, MS1 factor: 0.8, Gap filling by compulsion: True.

**Evaluation of feature detection.** The detection of a feature requires matching the m/z values within 5 ppm and retention time within 6 seconds, but does not require unique matching. The verified 402 features in Figure 4A were based on three samples (batch4_MT_20210729_003G, batch4_MT_20210729_003C, batch4_MT_20210729_003K). Data were processed using Thermo Scientific Compound Discoverer (v3.3), and a list of 402 manually verified features were created by visual inspection in FreeStyle (v1.8 SP2). The manually certified features in Figure 4B were given in the original publication (Chen et al, 2021), in the three unlabeled yeast samples with negative ionization. The true features in the NIST SRM 1950 sample were manually verified, and the list of 39 m/z features is given in the GitHub repository (**Data availability**). For the credentialed E. coli data (SZ22), the two feature tables were annotated separately using khipu (Li et al, 2023) to identify isotopic patterns. A pattern is considered valid when at least one M0 ion and one ion with more than one 13C labels are present in the compound group. The valid groups were combined from XCMS and asari, and [M+H]+ ions were compiled into the 643 “certified” features.

**Evaluation of computational performance.** The evaluation of computational performance was performed on a desktop computer with Intel i7-8809G CPU and 32 GB of memory, running Mint Linux 20.2. The asari version was 1.10.6. The XCMS version was 3.18.0. The R script for XCMS is provided in asari repository (https://github.com/shuzhao-li/asari) under doc/ directory. The time and memory use was measured by `/usr/bin/time –v`, and “User time” was used as CPU time (equivalent to CPU time used on a single core).

**Data structures in asari.** Examples of key data structures are given here as Python dictionaries (JSON compatible):

```
sample: {
    ‘input_file’: '',
    ‘list_scan_numbers’: [],
    ‘list_retention_time’: [],
    ‘rt_cal_dict’: {},
```

```
        'list_mass_tracks': [],
        }

    mass_track: {
        'id_number': 999,
        'mz': 230.01808166503906,
        'intensity': array([
            19265,  23414,  19809,  24195,  27025,  32111,  37948,  17387,
            33759,  32037,  24933,  15248,  30890,  24147,  37816,  52205,
            19562,  44433,  40049,  40032,  66720,  37805,  41621,  51155,
            52528,  41309,  51672,  36679,  63472,  55880,  63660,  68534,
            61723,  68821,  39000,      0,      0,      0,      0,      0,
                0,  59430,  91510, 220216, 254759, 273220,  64931,  25042,
                0,      0,  18063,  16858,  37267,  34924,  35216,  44651,
            36903,  51591,  31158,  31455,      0,      0,  58454,  48568,
            47294,  37701,  59255,  45277,  32742,  57048,  59220,  55310,
            71600,  51937,  52947,  54520,  46489,  56674,  62587,  55151,
            61771,  76765,  7699, ...,
                0,      0,      0,      0,      0,      0,      0,  10539,
                0,      0,      0,      0,      0,      0,   8404,      0,
                0,      0,      0,      0,      0,      0,      0,      0,
                0,   8164,      0,      0,      0,      0,      0])
    }

    feature:{
            "id_number": "F3943",
            "parent_masstrack_id": 2791,
            "mz": 313.23848724365234,
            "apex": 156,
            "left_base": 147,
            "right_base": 168,
            "rtime": 150.344,
            "rtime_left_base": 145.302,
            "rtime_right_base": 157.107,
            "peak_area": 153629884,
            "height": 18384026,
            "representative_intensity": 153629884,
            "snr": 217,
            "goodness_fitting": 0.9769980444055228,
            "cSelectivity": 1.0,
        },

    empirical_compound: {
            "interim_id": "kp480_314.2458",
            "neutral_formula_mass": 314.245709575,
            "neutral_formula": "C18H34O4",
            "Database_referred": [],
            "identity": [],
```

```
"MS1_pseudo_Spectra": [
{
    "apex": 156,
    "peak_area": 153629884,
    "height": 18384026,
    "left_base": 147,
    "right_base": 168,
    "goodness_fitting": 0.9769980444055228,
    "cSelectivity": 1.0,
    "parent_masstrack_id": 2791,
    "mz": 313.23848724365234,
    "snr": 217,
    "id_number": "F3943",
    "rtime": 150.344,
    "rtime_left_base": 145.302,
    "rtime_right_base": 157.107,
    "representative_intensity": 153629884,
    "id": "F3943",
    "isotope": "M0",
    "modification": "M-H-",
    "ion_relation": "M0,M-H-",
    "parent_epd_id": "kp480_314.2458"
},
{
    "apex": 156,
    "peak_area": 26869245,
    "height": 3362120,
    "left_base": 150,
    "right_base": 168,
    "goodness_fitting": 0.9598939375019998,
    "cSelectivity": 1.0,
    "parent_masstrack_id": 2805,
    "mz": 314.2418899536133,
    "snr": 332,
    "id_number": "F4045",
    "rtime": 150.344,
    "rtime_left_base": 146.982,
    "rtime_right_base": 157.107,
    "representative_intensity": 26869245,
    "id": "F4045",
    "isotope": "13C/12C",
    "modification": "M-H-",
    "ion_relation": "13C/12C,M-H-",
    "parent_epd_id": "kp480_314.2458"
},
{
    "apex": 161,
    "peak_area": 10973710,
```

```
                "height": 1116800,
                "left_base": 155,
                "right_base": 173,
                "goodness_fitting": 0.588704815611131,
                "cSelectivity": 1.0,
                "parent_masstrack_id": 2616,
                "mz": 295.2277374267578,
                "snr": 46,
                "id_number": "F4216",
                "rtime": 153.157,
                "rtime_left_base": 149.783,
                "rtime_right_base": 159.93,
                "representative_intensity": 10973710,
                "id": "F4216",
                "ion_relation": "M-H2O-H[-]"
            }
            ],
            "MS2_Spectra": [],
            "list_matches": [
            [
                "C18H34O4_314.24571",
                "M-H[-]",
                2
            ],
            [
                "C18H36O5_332.256274",
                "M-H2O-H[-]",
                1
            ]
            ]
        }
```

### Supplemental Tables

**Table S1. Pairwise, unambiguously matched features between processing tools on the HZV029q dataset.** MZmine is using the ADAP wavelets algorithm for elution peak detection, and MZmine(L) is based on the local minimum search algorithm. Processing parameters and software versions are given in the Methods section. In the comparison between two result tables, a feature is considered matched when the m/z difference is within 5 ppm and retention time is within 6 seconds; if multiple matches are found within the parameters, the pair with closest retention time is chosen. Alternative method of choosing among multiple matches by closest m/z values produced similar results.

| HZV029q (184 files of repeated QC sample, positive ionization) | | | | | |
|---|---|---|---|---|---|
| *XCMS* | *MZmine* | *MZmine(L)* | *MS-DIAL* | *asari* | |
| 10901 | 6186 | 6227 | 5421 | 5360 | *XCMS* |
| | 42099 | 17223 | 14286 | 13164 | *MZmine* |
| | | 24837 | 12527 | 11963 | *Mzmine(L)* |
| | | | 54863 | 10624 | *MS-DIAL* |
| | | | | 22440 | *asari* |

**Table S2. Pairwise, unambiguously matched features between processing tools on the Yeast2021 dataset.** Methods are the same as in Table S1.

| Yeast2021 (3 files, yeast culture extracts, negative ionization) | | | | | |
|---|---|---|---|---|---|
| *XCMS* | *MZmine* | *Mzmine(L)* | *MS-DIAL* | *asari* | |
| 6043 | 4728 | 4364 | 2609 | 3013 | *XCMS* |
| | 11290 | 6498 | 2939 | 3435 | *MZmine* |
| | | 18153 | 2760 | 3395 | *Mzmine(L)* |
| | | | 4166 | 3108 | *MS-DIAL* |
| | | | | 5341 | *asari* |

**Supplemental Figures**

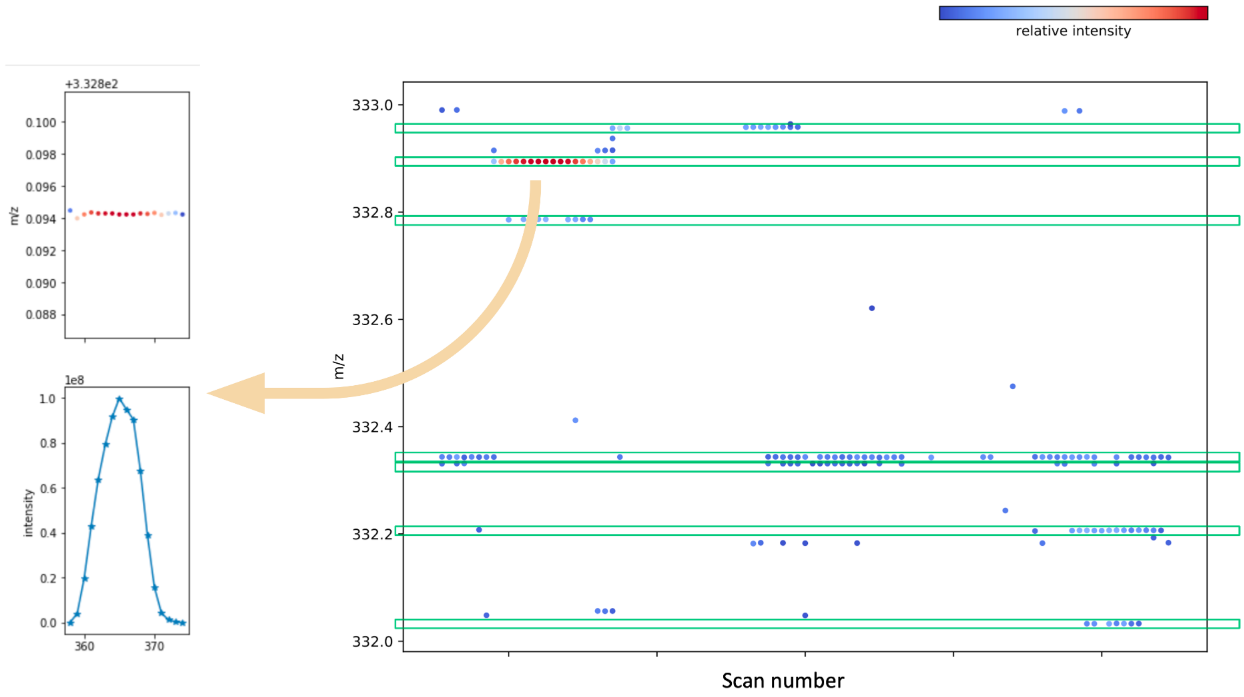


**Figure S1 Illustration of mass tracks in a single sample.** The region has 7 mass tracks marked by green boxes spanning horizontally, each of a unique m/z value. A peak is detected from the track indicated by the yellow arrow.

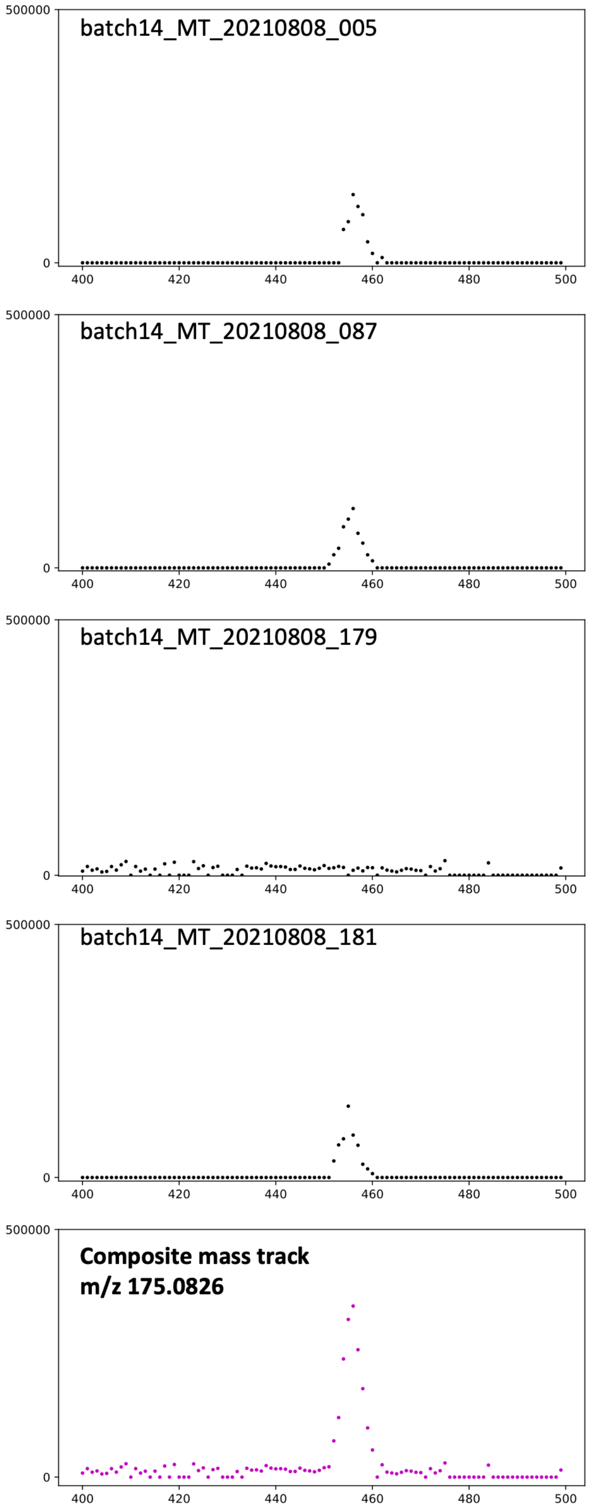


**Figure S2. Composite mass track enhances peak detection by aggregating signals from all samples.** An example is shown at m/z 175.0826 in the MT02 dataset. Four individual samples are shown on top, and the composite mass track at bottom.

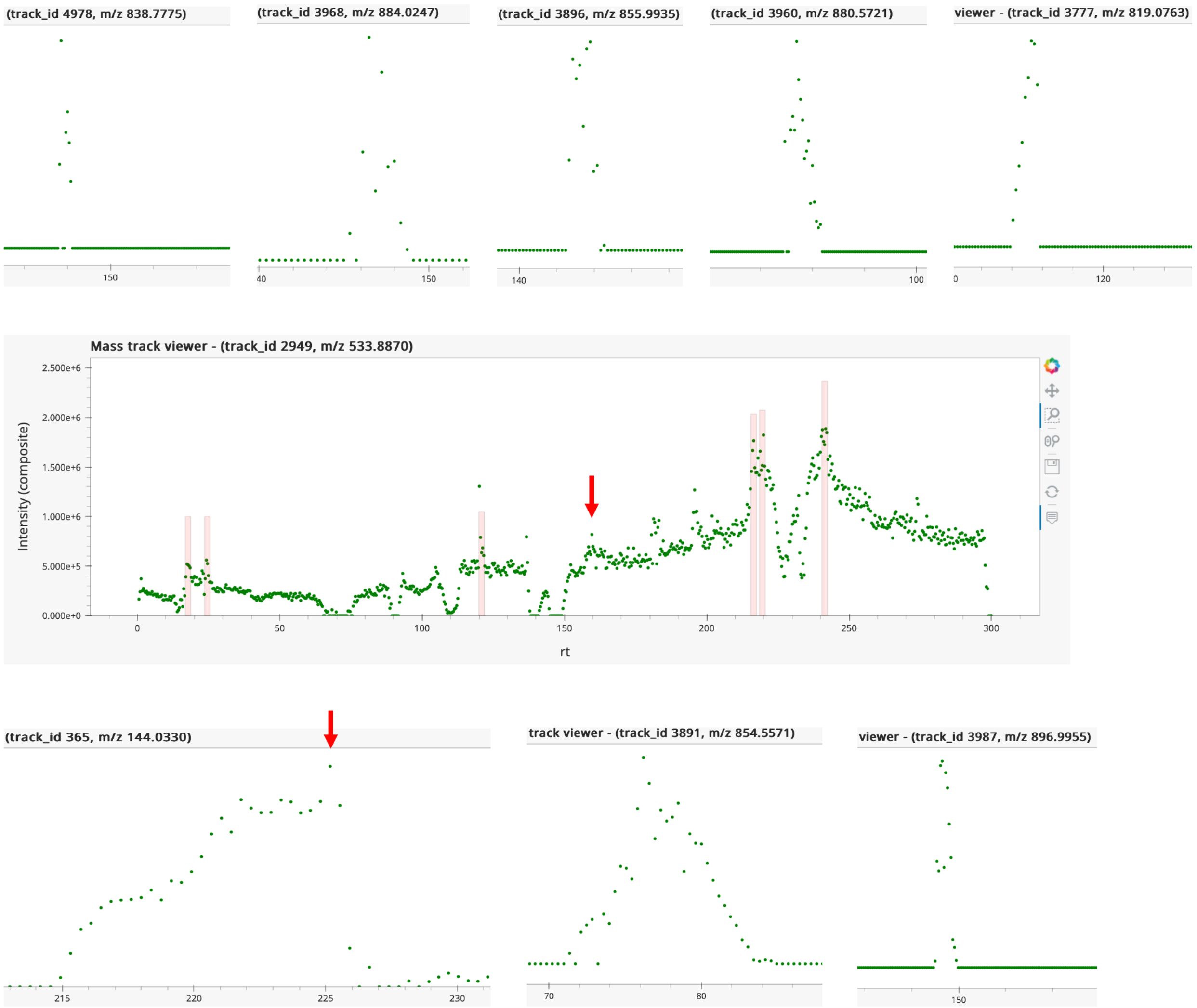


**Figure S3. Manually verified features that are not reported by asari in the HZV029 dataset (corresponding to Figure 4A).** Part or full mass track is shown. The red arrow indicates the missed features in complex tracks.

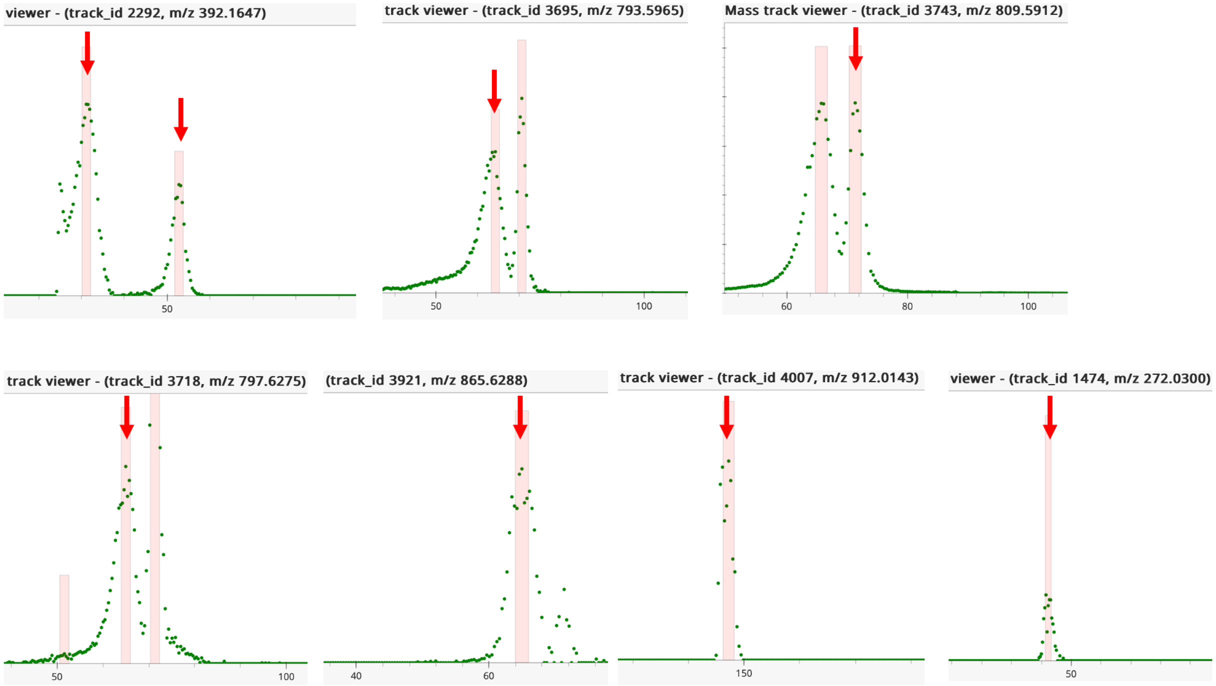


**Figure S4. Manually verified features that are not reported by MS-DIAL in the HZV029 dataset (corresponding to Figure 4A).** Partial mass tracks are shown. The red arrow indicates the missed features. These features were all detected by asari.

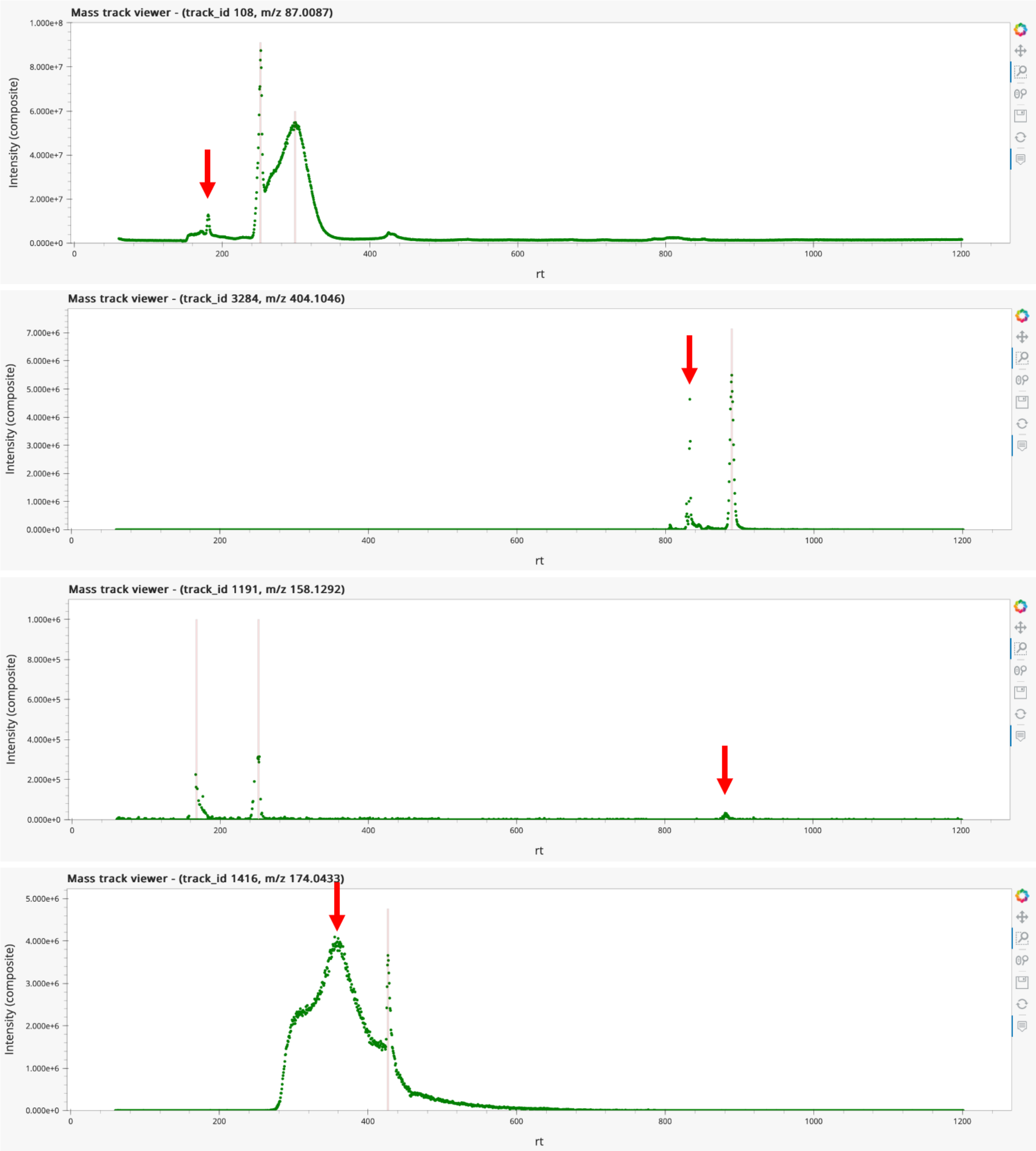


**Figure S5. Four manually verified features are not reported by asari in the Yeast2021 dataset (corresponding to Figure 4B)**. The top two were due to too few valid data points in a peak. The third case was caused by peak height below threshold, and detectable by lowering the threshold. The bottom case was due to high local noise.

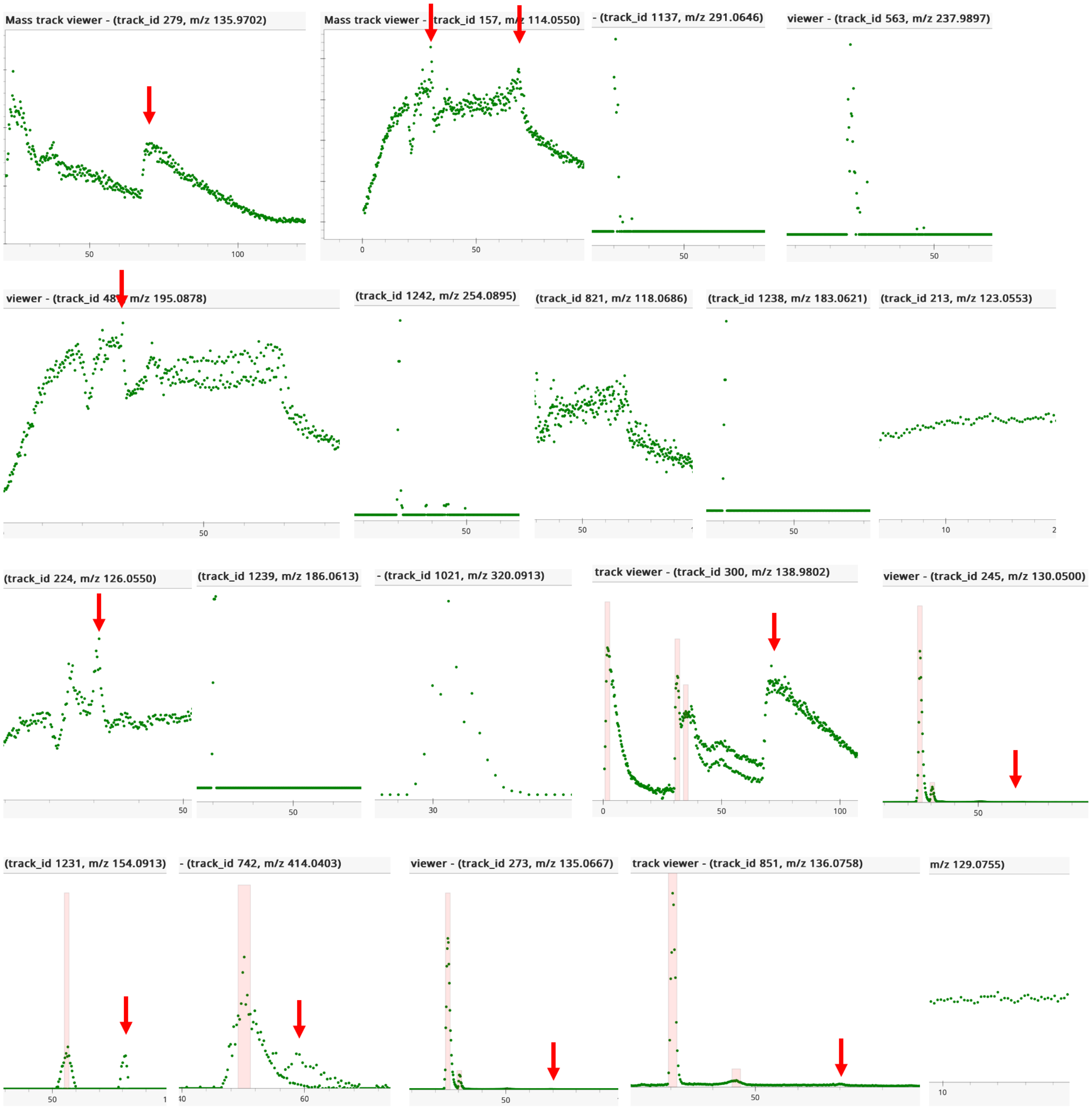


**Figure S6. Features not reported by asari in the SZ22 E. coli dataset (corresponding to Figure 4C).**

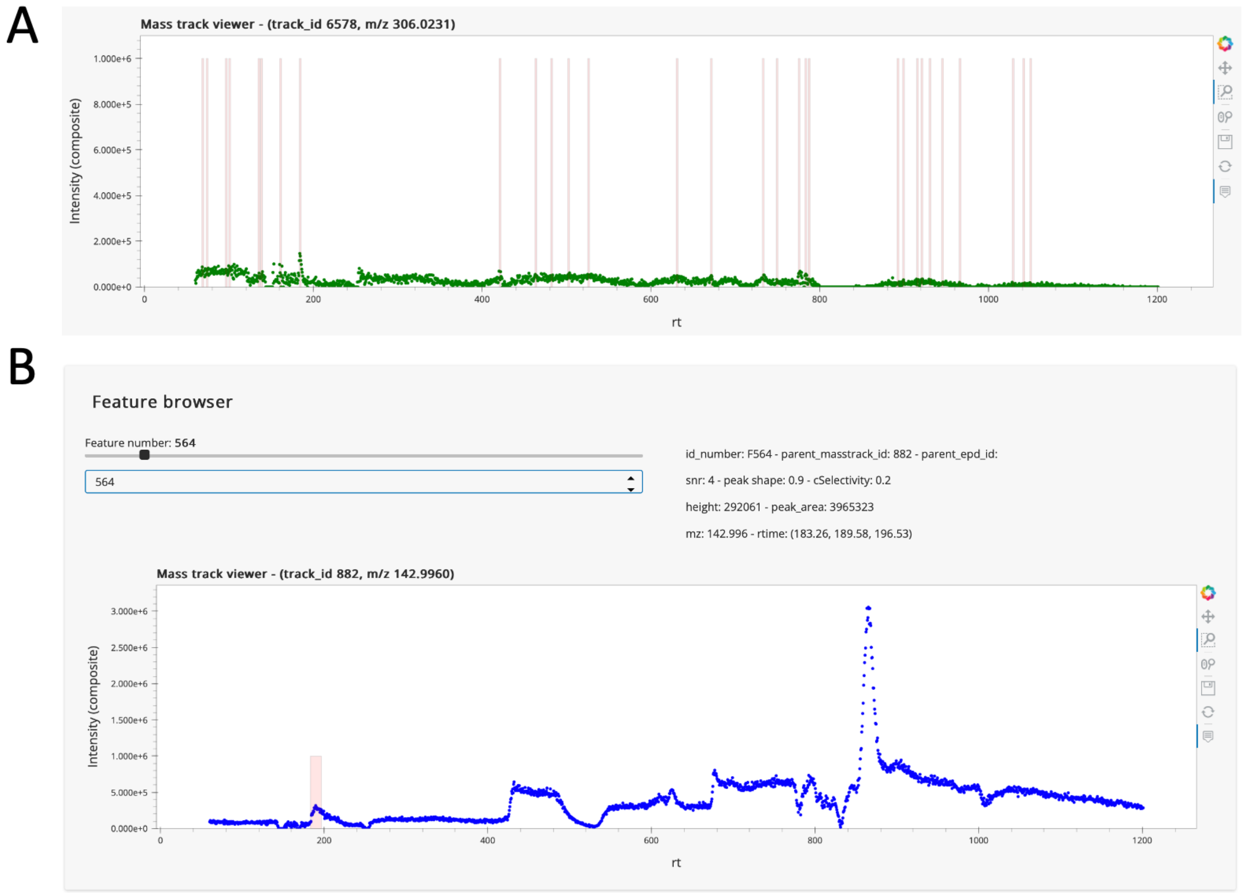


**Figure S7. Example chromatograms for illustrating problems in peak detection.**

A) Inappropriate parameters can lead to many low-quality peaks. Each pink vertical line indicates a peak.

B) The chromatography shows higher baseline in later elution time. This is from Feature Browser in asari dashboard, which shows only the current not all features on the mass track.
